# Efficacy of bacteriophage therapy against vancomycin-resistant *Enterococcus feacalis* in induced and non-induced diabetic mice

**DOI:** 10.1101/2021.01.21.427594

**Authors:** Ajay Kumar Oli, Nagveni Shivshetty, Likyat Ahmed, Manjunath Chavadi, Rahul N Kambar, R Kelmani Chandrakanth

## Abstract

The antibiotic resistance of an organism has become an endeavor worldwide. The resistance of bacteria is impacting the immunocompromised patient, especially with diabetic. The bacteriophages are more specific in the elimination of the infections caused by the organism. One of the significant bacteria is Vancomycin-resistant *Enterococcus faecalis* (VREF), emerging as bioburden rendering infectious diseases in developing countries. We attempted to treat infection of VREF in the induced and non-induced diabetic mice model along with antibiotic and phage treatment. The phage has shown more efficiency than vancomycin antibiotic alone and combined therapy with antibiotic phage treated. The considering phage therapy an alternative source for the treatment in the insulin-dependent diabetic with an infection of bacteria.

## Introduction

*Enterococcus feacalis,* a gram-positive facultative anaerobic bacterium, is mainly present in humans and animals’ microbiota. It has now emerged as one of the significant causes of human clinical infections. Antibiotics are the ‘miracle drugs used for treating bacterial infections. Due to the cheaper availability, these are commonly used and widely spread worldwide, leading to irrational uncontrolled uses of antibiotics, causing bacterial resistance**[1,2]**. Antimicrobial resistance is threatening to human health. It is one of the most significant challenges of our civilization. It could be possible to cause an estimated 10 million mortalities per year by 2050 **[3]**. Glycopeptide antibiotic-resistant enterococci have become a significant threat to hospitalized patients causing breaks that increase morbidity, mortality, and health care **[4].** Enterococci can acquire resistance to multiple antimicrobial agents rapidly totheir pathogen via transposons and plasmids **[5, 6, 7]**. The emergence of VRE is becoming a grave public health concern, especially in developing countries where there is a lack of availability of effective antibiotics **[8].** The natural predators of bacteria are bacterial viruses known as bacteriophages or phages. These are ubiquitously found and are estimated to be present at numbers equivalent to a trillion per grain of sand on earth **[9].** The bacteriophages can be an alternative potential antibacterial therapeutic agents against multidrug-resistant pathogens **[10]**. Phage therapy is rapidly evolving in medicinal uses in life-saving and undergoing multiple clinical trials. The present study checks bacteriophage therapy’s efficacy against vancomycin-resistant *Enterococcus feacalis* in diabetic and non-diabetic induced mice models.

## Material and Methods

### Isolation of Bacteriophages

*E. faecalis* specific bacteriophage was isolated by a regular enrichment procedure consistent with Sheng **[11]** from raw sewage collected from a municipal sewage treatment system. Briefly, 50-ml sewage samples were centrifuged at 5,000 rpm for 10 min at 4°C. The supernatants were filtered through a 0.45μm-pore-size filter (Pall Life Sciences), 10 ml filtrate was added to 1 ml 10 X LB broth, and 100μl (~ 10^8^ CFU) of each *E.faecalisstrain* was added. The mixture was incubated at 37°C for 18 h with aeration (180 rpm). Debris and bacteria were removed by centrifugation at 10,000 rpm for 10 min, and supernatants were filtered through a 0.2μm-pore size filter (Pall Life Sciences). Phage activity within the supernatant was tested by a spot assay that entailed placing 5μl of the supernatant on BHI agar seeded with *E. faecalis* strains. The plates were evaluated for plaques formation after 18 h at 37°C. Lysispositive supernatants were serially diluted, and plaques were isolated and purified, employing a soft agar technique.

### Broth Culture of Phage

*E. faecalis* 122 was inoculated 1:100 ratio of 200 ml of BHI broth and incubated at 37°C in a shaking incubator (240 rpm). As the optical density OD at 600 nm showed 0.1 reading bacteriophage with 10^7^ plaque-forming unit (PFU)/ml, is then added to the cultured bacterial flask. Every 15 min, samples were collected from the broth, and OD readings of infected and uninfected cells were measured at 600 nm. Completely lysed cultures were then centrifuged (10,000 rpm, 5 min, 20°C), and chloroform-treated phage was titrated within the supernatant.

### Adsorption rate, latent period, and phage burst size

The procedure was carried according to the previous study by Adams **[12]**. Briefly, to check live the adsorption rate, 1 ml phage (1.5 x 10^3^ PFU/ml) and 1 ml VREF122 (5 x 10^8^ CFU/ml) were mixed, and the number of free phage particles was resolute after treatment with chloroform (200μl). To confirm the latent period and burst size, VREFbacteria (5 x 10^8^ CFU/ml) were incubated with phage (3 x 10 PFU/ml) for five mins, washed with cold LB broth to get rid of free phage particles, and so resuspended in fresh medium. The cell suspension was periodically titrated for newly produced phage on the VREF122 culture lawn.

### Phage purification and Storage

Theisolated colony of VREF122 was inoculated into BHI broth. Culture with OD of 0.1 is used to infect with phage GACP 2×10^7^ PFU; the culture was co-cultivated for 18 h 37°C in a shaking incubator (240 rpm). Polyethylene glycol-8000(PEG) or NaCl was added to the lysate to a final concentration of 20% or 0.5 M respectively and incubated for 1 h at 4°C. After centrifugation at 10,000 rpm (16 min at 4 °C) in the centrifuge, polyethylene glycol (PEG 8000) was added to the supernatant to a final concentration of 10%. The lysate was incubated overnight at 4°C with gentle stirring. Polyethylene glycol-precipitated phages were collected by centrifugation at 15,000 rpm for 20 min. The resulting pellets were suspended in 3 ml of phage buffer (20mM Tris-HCL [pH 7.4], 100mM NaCl, 10 mM MgSo_4_), filtered through 0.2μm bacterial filters, and phage filtrate was recovered and dialyzed against phage buffer. Purified phage GACP was stored in aliquots of phage buffer at −20°C **[13].**

### Electron Microscopy of Phage GACP

A drop of the phage GACP suspension is applied to a formvar carbon-coated copper grid for five mins; the rest was removed with a pipette and negatively stained with 2% uranyl acetate (TAAB Laboratory UK). After 10 min, the liquid is removed withpaper. The grids were examined in an exceedingly TechaniBiotwin (Philips), transmission microscope, (Netherland) at 80 Kv (Magnification, X105, 000 or X160, 000) **[14].**

## Animals Experiments

### Animals

Specific pathogen-free, colony bred, virgin adult Swiss mice (Wistar strain) of both sexes weighing 25±0.8 grams were obtained from the experimental animal facility of Luqman College of pharmacy. Animals were used with the Institute Animals Ethics Committee (Reg.No. 346/CPSCEA). Animals are fed with pellets (VRK Nutrition Solutions, Sangli, Maharashtra, India Ltd.,) and water adlibitum. After randomization into groups and experiment initiation, the mice were acclimatized for seven days under standard laboratory conditions of temperature 24-28°C, ratio humidity of 60-70%, and 12 h day and night cycles). Mice were randomly assigned to groups were housed (in Luqman Pharmacy College’s animal house at Kalaburagi) in cages of two or three until experiments started.

### Preparation of Streptozotocin (STZ) doses

Mice were made diabetic by consequent intraperitoneal injections of various concentrations of STZ (Hi media, Mumbai). STZ was first weighed individually for every animal per the weight and so suspended in 0.2 ml citrate buffer (0.1M Citrate buffer, pH 4.5) just before injection.

### STZ-induced Diabetes

To induce diabetes, the mice were treated with STZ [60, 120, and 180 mg (Kg body weight)^−1^ in 0.1 M Citrate Buffer, pH 4.5] by intraperitoneal administration for 3 consecutive days. At the time of the STZ injection, the mice are fasted for 14-16 h but had free access to water. After the STZ treatment, all the mice were returned to their cages and given free access to food and water. Blood sugar levels are monitored after the stabilization period of ten days **[15].**

### Collection of blood from Experimental mice

The mice are weighed weekly, and blood glucose levels are determined to assess hyperglycemia employing a handheld glucometer. Blood samples were collected from the tail vein, and also the fasting blood sugar levels were estimated using an Acu-Check Active (Indiapoils) electronic glucometer.

### Pathogen

The pathogenic strain of VREF 122 (Catalase negative, MDR, high-level resistant vancomycin, mouse virulent) was employed in all experiments. This organism was grown in BHI broth was to prepare bacterial inocula. To induce infection in mice, 10^6^-10^9^ CFU/ml count of *E. faecalis* washed in 0.2ml of 0.9% PBS was used.

### Experimental induction of VREF122 bacteremia in Diabetic and non-diabetic mice

For each infection experiment, 6 to 8-week old diabetic and non-diabetic mice were divided into two groups, and each group was given inocula of varied sizes. Preparation of the infecting bacteria was as follows, VREF122 isolate was grown in the BHI broth medium at 37°C and centrifuged at 8,000 rpm for five mins. The cell pellet was washed with normal PBS, centrifuged again under identical conditions, and at last resuspended in 10 ml PBS. After appropriate dilution, turbidity at 600nm is measured to see bacterial cell numbers. To evaluate minimum lethal dose (MLD), serial dilutions of VREF122 were injected intraperitoneally (i.p.) into diabetic and non-diabetic mice in 0.1ml aliquots. After infection, mice are kept under standard laboratory conditions with free access to food and water. Twelve mice were used for each dose; non-diabetic and diabetic mice’s survivals were then measured two days postinfection **[16]**. Mice inoculated with VREF122 were observed for their state of bacteremia. They were scored for their state of health on anarbitrary scale of 5 to 0, supported by progressive disease state reflected by several clinical signs. An unremarkable condition is scored as 5, slight illness, defined as lethargy and ruffled fur, is scored as 4, moderate disease, characterized as severe lethargy, ruffled fur, and hunched back, is scored as 3; severe illness, with the above signs plus exudative accumulation around partially closed eyes, is scored as 2, a moribund state is scored as 1, and death is scored as 0. The dose which shows 50% lethality is taken because of the optimum LD_50_ dose.

### Detection of Pathogen in Circulatory System

An experiment to determine the entry of VREF122 infection in mice from the peritoneal cavity into the circulatory system is executed. The bacteria were injected i.p. followed by the gathering of blood samples from the tail vein, and approximately 0.1ml was collected into a 1.5 ml E-tube containing 10μl EDTA (100mM). The blood samples obtained were cultured on bile esculinazide agar.

### Vancomycin Treatment in VREF122infected mice

The animals were divided into four groups, two of diabetic and non-diabetic, followed by induction of VREF122 strain. One group was kept as the control without antibiotic induction, and the remaining were injected with the antibiotic of vancomycin [2mg(Kg body weight)]^−1^ after 20 min of pathogen administration. Further, the antibiotic was routinely induced for five days.**[17].**

### Efficacy of phage GACP in bacteremia mice model

The phage’s efficacy was studied in six groups of mice (Twelve mice in each group) and was challenged by the intraperitoneal injection of VREF122. Each group was treated with a single injection of GACP at 3×10^10^ PFU after 20 min of bacterial induction. The experiment is observed for 30 days.

### Combined therapy in bacteremia mice model

The combined therapeutic potential is determined usingvancomycin antibiotic and phage GACP in diabetic and non-diabetic VREF122bacteremia mice model. The effect of antibiotic and phage dose is studied within the six groups of mice (Twelve mice in each group). A single dose of Vancomycin antibiotic was administered i.p after 20 min bacterial induction and followed by phage with a dose of 3×10^11^ PFU.

### Quantification of the immunological reaction of phage GACP in mice

Bacteriophages are beneficial agents for studies of humoral immunity. They are potent antigens that cause no toxic effect on the health of humans. **[18, 19].**

During the experiment of mice were treated with phage GACP (3×10^10^ PFU) through i.p injection. At various time points, mice blood was collected from the optic vein and cultured. Subsequently, the collected blood was subjected to indirect Enzyme-Linked Immunosorbent Assay to detect antibodytiters of IgG IgM antibody titers in sera of diabetic and non-diabetic mice as described **[20]**.

## Results

### Bacteriophage isolation

The MDR strains of *E.faecalis* were inoculated with sewage samples obtained from different local drainages. And supplements were harvested, and it is subjected to the spot test. The spot test for phage formed a clear zone on a specific number of the *E.faecalis* isolates. Among them, VREF 122 isolate tested manifested turbid zone is shown in figure results indicates that the phage possessed a lytic nature against an isolate of VREF 122

The adsorption rate, latent period, and burst size of phage thought to be biological parameters for phage infection measurements are shown in the figure. Comparing these parameters of ϕGACP, the adsorption rate within 5 min was about 90%, the latent period was about 25min, the burst size was about 110–120 PFU/cell. Medians of latent period and burst size in tailed phage were typically 40–60min and 50–100 PFU/cell, respectively, and were capable of infecting a large spectrum of *E. faecalis* strains, causing completed and confluent lysis. Thus, it was indicated that ϕGACP as a therapeutic agent to manage infections caused by *MDRE.faecalis,* as shown in figure 2.

**Figure 1.**
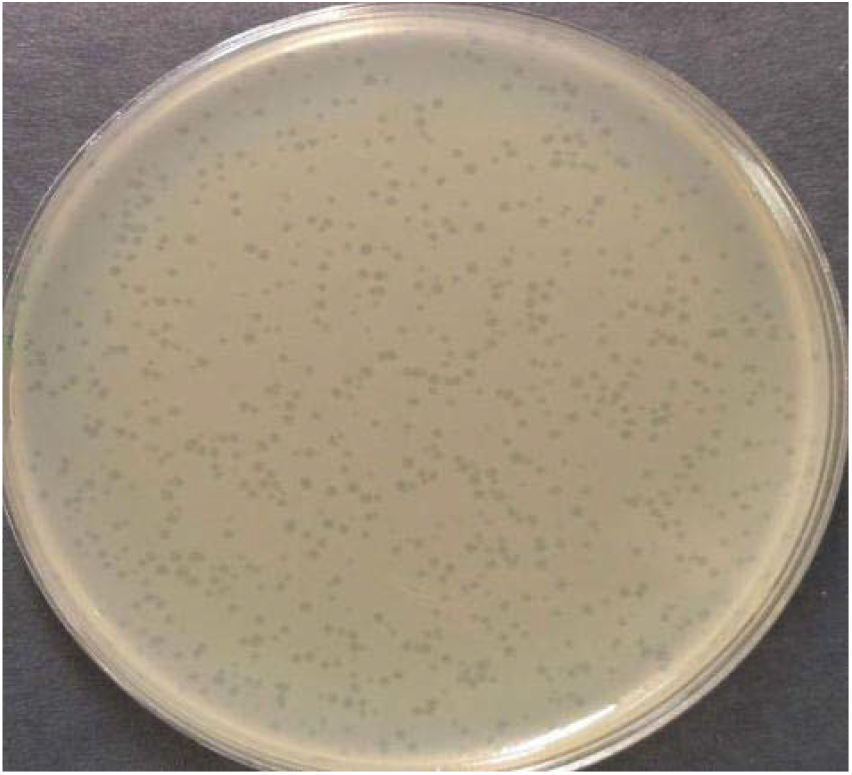
Plate showing plaque formation of bacteriophage on BHI agar media against VREF122.

**Figure 2.**
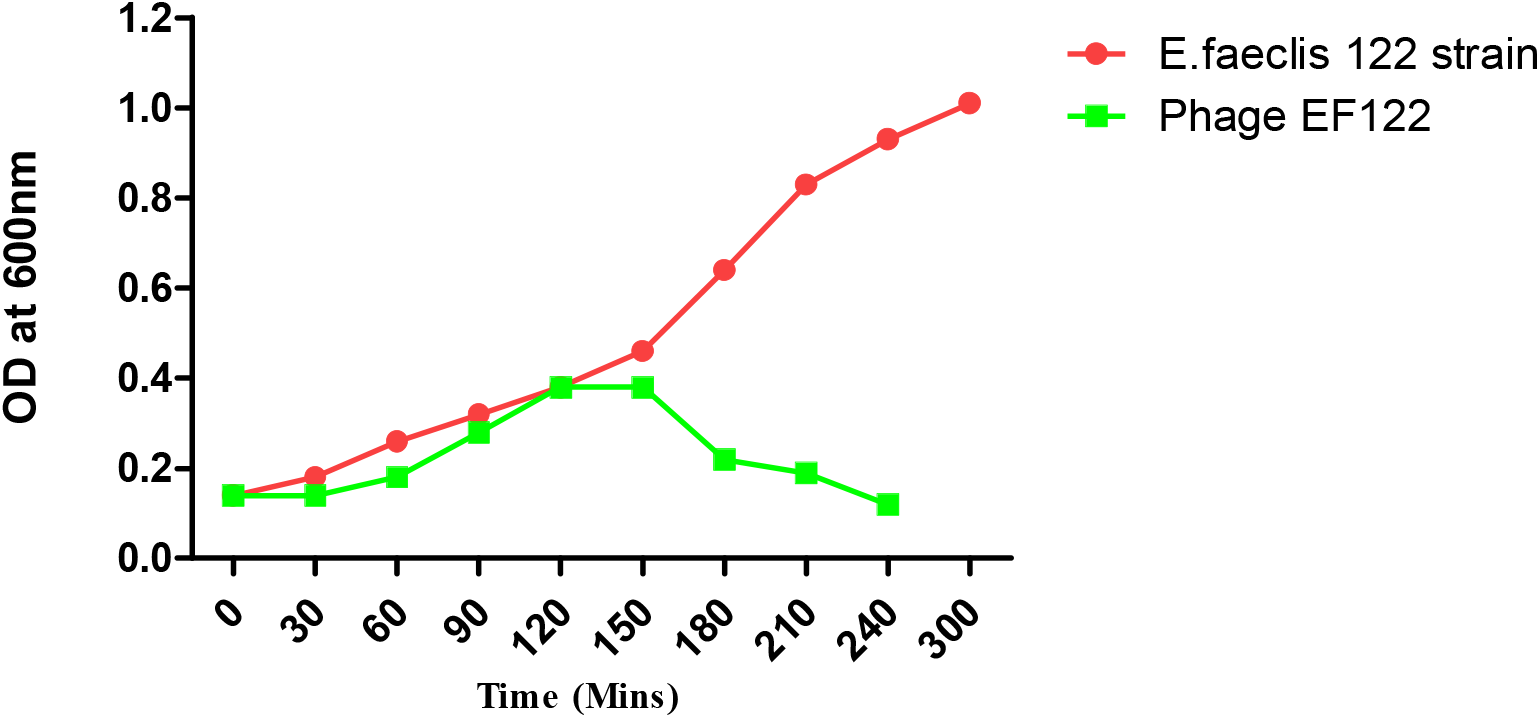
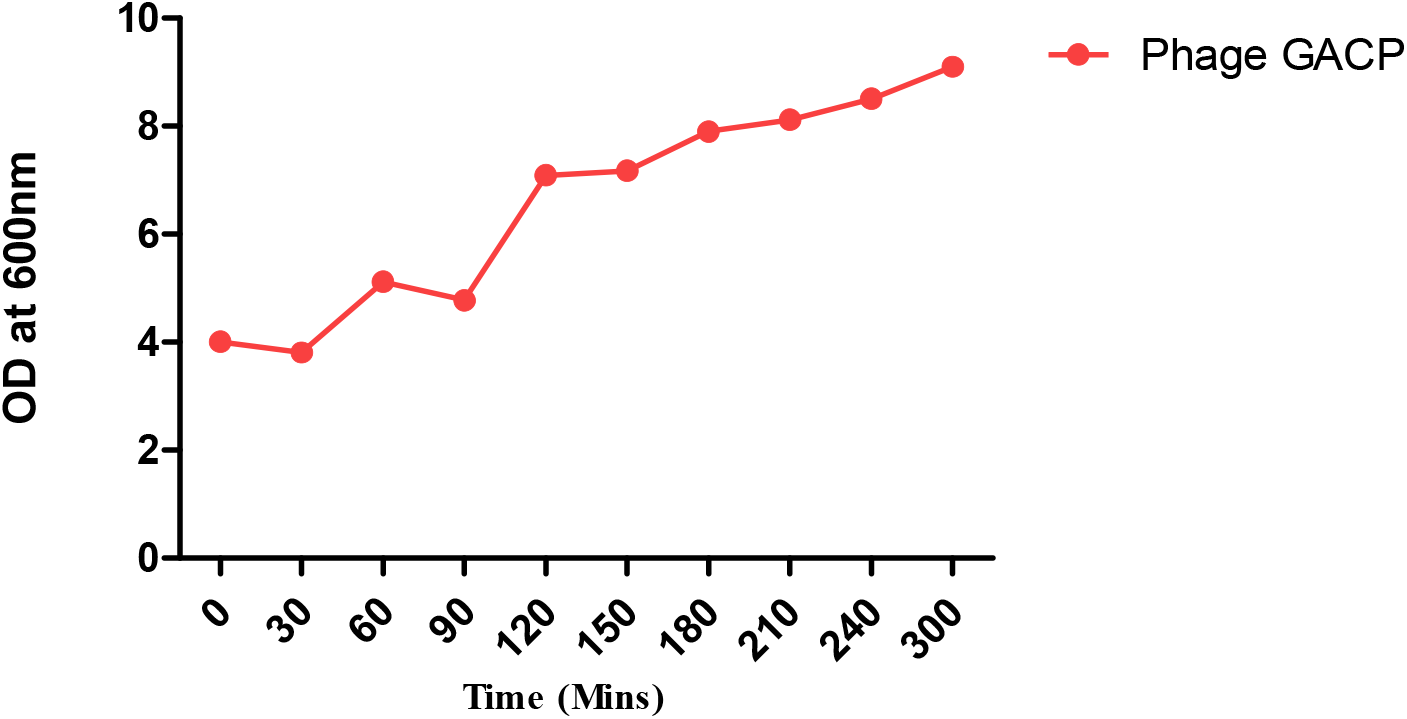
Lysis *of Enterococcus faecalis* EF122 isolate by the phage in broth culture. A. OD development of an uninfected control culture (VREF122) and parallel Culture infected with Phage GACP B. Progeny phage release from the phage infected culture depicted in fig A. Phage infectivity was measured by plaque assay.

### Transmission electron microscope

Phage GACP possesses isometric heads of 100 nm in diameter and conspicuous capsomers, striated 140 long tails, a double base plate, and globular structures at the tail tip, as shown in Figure.3. It belongs to thefamily myoviridae, subfamily Spounavirine family.The proposed subfamily contains the ICTV recognized genus “SPO1-like viruses.”

**Figure 3:**
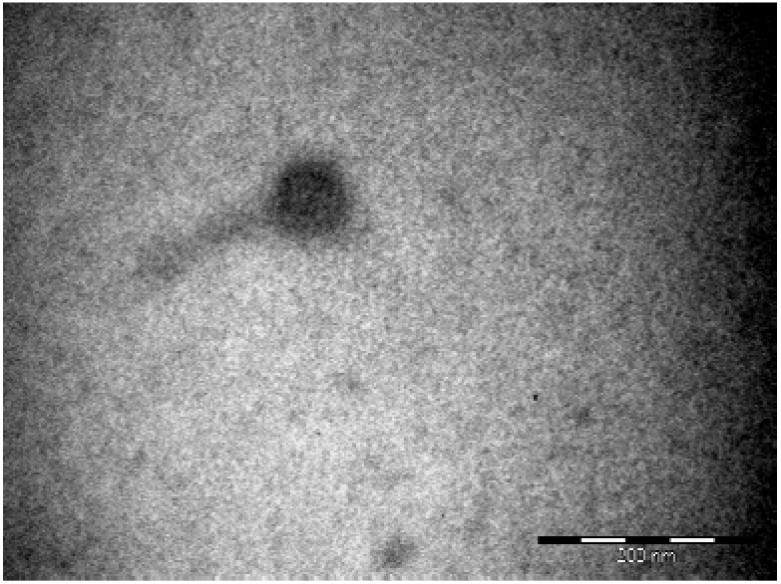
TEM of phage GACP magnification X68, 000.

### Streptozotocin (STZ) - induced diabetes in Experimental Mice

Mice induced with multiple doses of STZ [150 mg (Kg body weight)-^1^] have shown chronic hyperglycemia as represented in Table.1. One group received a dose of 180 mg of STZ (Kg body weight)-^1^ died after 20 days. Groups of mice that received a concentration of 120 mg of STZ (Kg body weight)-^1^ were healthy and alive for two months. Although the diabetic mice exhibited glycosuria, they also developed ruffled hair, appeared thinner, and consumed more food per day than the untreated group.The glucose level ≥of 300 mg/dl was considered as diabetic mice.

**Table.No.1.** Fasting Blood Glucose Concentration in mice (n=12) administered with various doses of Streptozotocin

| I.P doses of STZ (mg/Kg) | Fasting blood glucose (mg/dl) at day after STZ administration |  |  |
| --- | --- | --- | --- |
|  | Day 10 | Day 15 | Day 20 |
| 0 | 83 | 89 | 96 |
| 60 | 110 | 134 | 178 |
| 120 | 272 | 356 | 422 |
| 150 | 312 | 378 | 442 |
| 180 | 372 | 484 | 558 |

### Host screening of Vancomycin-resistant *E.faecalis* strain in diabetic and non-diabetic mice

A total of 12 VREF strain VREF 122 were selected based on its high resistant and multidrug-resistant nature and its susceptibility for phage GACP lytic multiplication. Finally, bacterial strain VREF 122 is set for induction of bacteremia in diabetic and non-diabetic mice.

### Induction of experimental bacteremia in diabetic and non-diabetic mice model with VREF 122 isolate

The lethal dose of *E. faecalis* to mice was resolute by injecting both diabetic and nondiabetic mice with varying cell concentration VREF122, starting from 3×10^6^ to 3×10^9^ cells /dose. Intraperitoneal injection of 3×10^6^ to 3×10^7^ didn’tdecrease the persistence rate of diabetic, whereas 10 to 20 percent of mice died during the next seven day observation period and 0 to 20 percent non-diabetic mice, respectively. In contrast, VREF122 3×10 cells showed a survival rate of 50 percent in diabetic and non-diabetic mice within 48 hrs. VREF 122, 3×10^8^ showed a 100% lethal effect within 48 h of injection in both groups of animals as shown in Figure 4 a and b; therefore, this dose 3×10^8^ CFU/ml was the optimal dose for the experiment.

**Figure 4:**
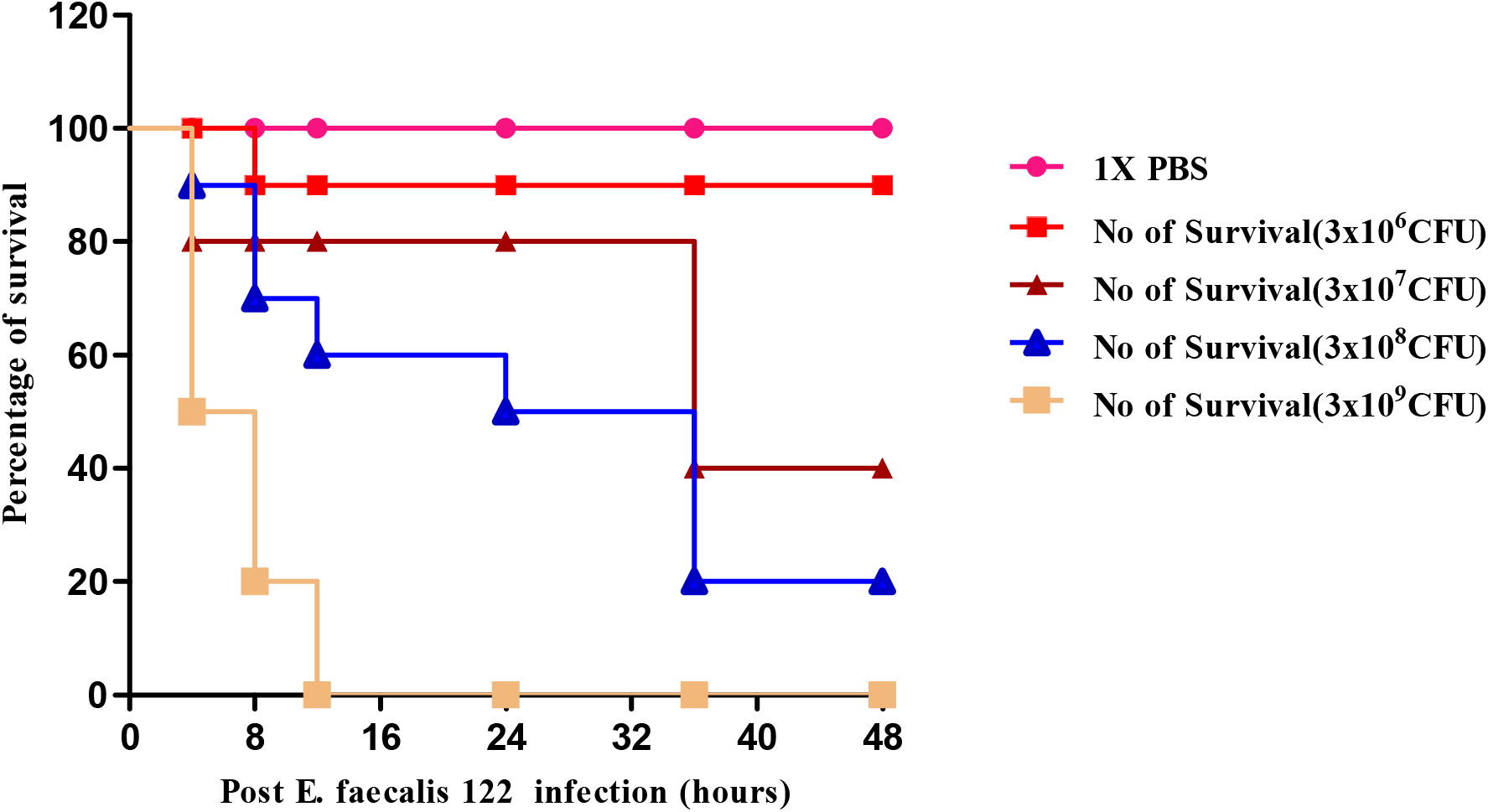
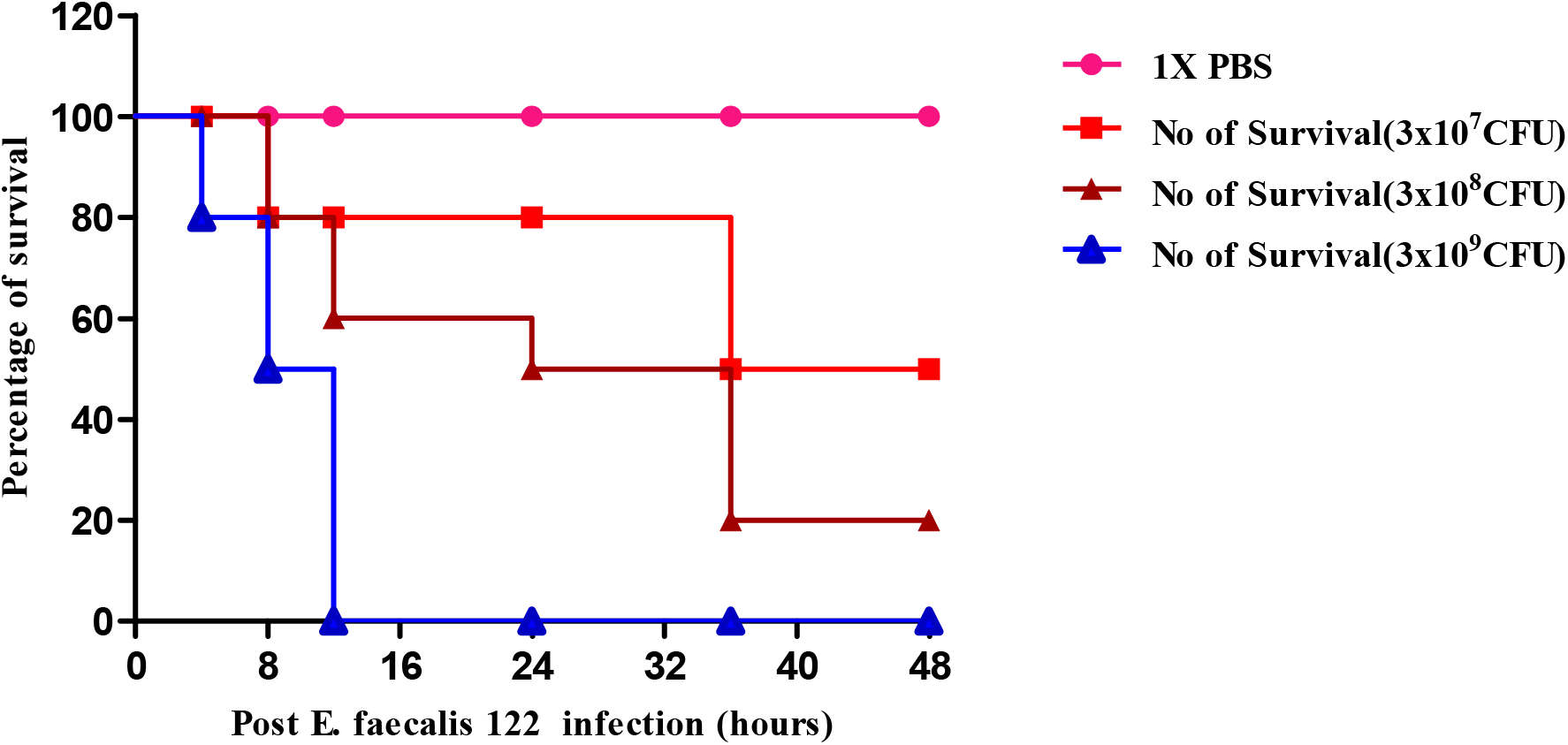
a. Determination of the Minimal Lethal Dose (MLD) of VREF122 in Diabetic mice 3×10^6^(A) 3×107 (B) 3×10^8^(C)3× 10^9^(D), 1X PBS Control CFU/ml represents percentage of the survival rates up to 48 hours after administration. b. Determination of the Minimal Lethal Dose (MLD) of VREF122 in Nondiabetic mice 1X PBS control,(A) 3×10^7^ (B)3×10^8^(C)3×10^9^(D) CFU/ml represents percentage of the survival rates up to 48 hours after administration.

### Protection of lethal bacteremic diabetic and non-diabetic mice by $ GACP

Different concentrations ofpurified phage GACP of 3×10^7^ to 3×10^10^ PFU were injected to check the protection activity in diabetic and non-diabetic mice from VREF 122 of lethal bacteremia. Figure 5a and b show that 3×10^10^ of $ GACP significantly rescued 100% diabetic and non-diabetic mice from fatal bacteremia. All live mice remained healthy for 30 days till the experiment was terminated. The phage dose effect on the health status of the infected animals was visible. Administration of a high dose 3×10^10^ of phage GACP alone to experimental mice also didn’t affect their soundness of survival of mice during the one month of observation.

**Figure 5:**
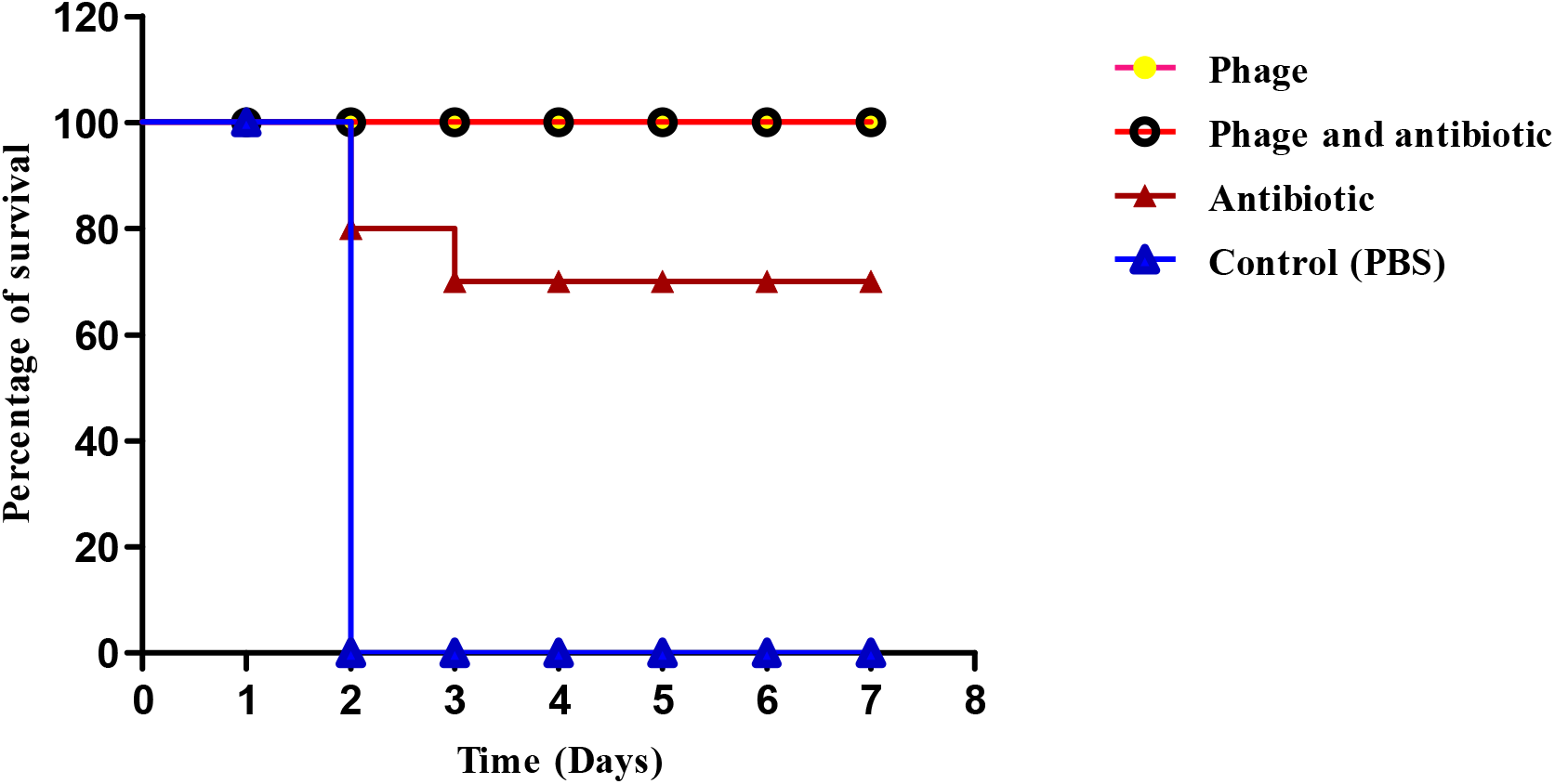
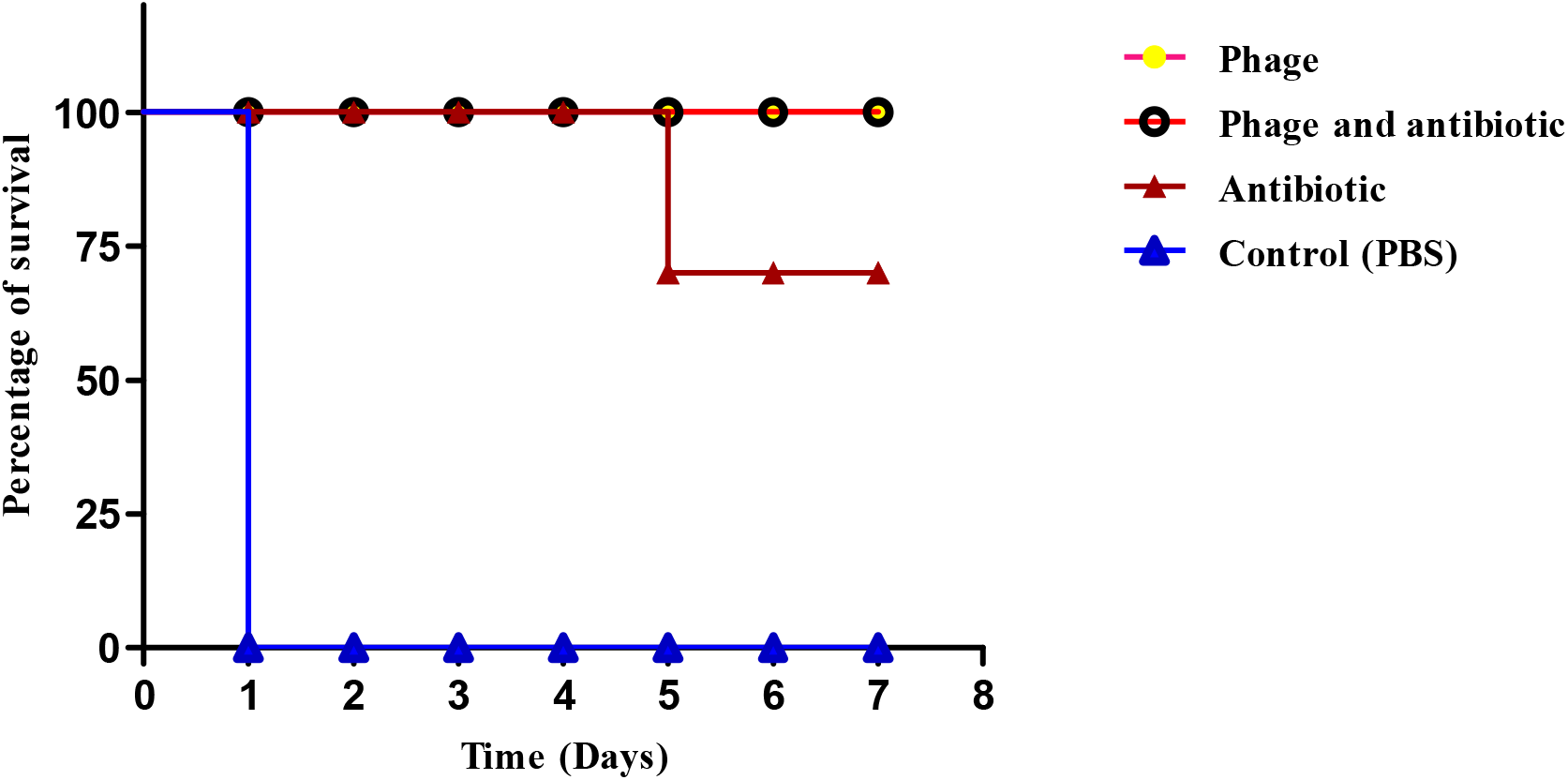
a Determination of the protection of VREF122 in Diabetic mice control PBS (A) 3×10^10^ single dose of phage (B)single dose of vancomycin antibiotic (C) 3× 10^10^ phage and single dose of antibiotic represents percentage of the survival rates up to 7 days after administration. b Determination of the protection of VREF122 in Non-Diabetic mice control PBS (A) 3×10^10^ single dose of phage (B) single dose of vancomycin antibiotic (C) 3× 10^10^ phage and single dose of antibiotic represents percentage of the survival rates up to 7 days after administration.

### Treatment efficacy of phage compared withvancomycin antibiotic in diabetic and nondiabetic mice

Phage’s effectiveness with vancomycin, antibiotic treatment of diabetic and non-diabetic mice, a single i.p. dose of vancomycin [2mg(Kg body weight)-^1^] after 20 min of VREF 122 results exhibited 70% protection in both diabetic and non diabetic mice from VREF122. Whereas, phage GACP of 3×10^10^ PFU has rescued 100% in both mice groups from VREF 122 bacteremia. There have been no survivors among untreated mice (in both groups) after two days of i.p injection 3×10^8^ CFU cells of VREF122 as shown in (Fig.5a & b). The protected mice were healthy for the next seven days of observation time.

### Protection of lethal bacteremia mice by combined treatment with GACP and Vancomycin antibiotic

A purified ϕ GACP of 3×10^7^ to 3×10^10^ PFU and a single dose of [2mg(Kg body weight)-^1^] vancomycin induced showed the protection efficiency of 80% and 90%in diabetic and nondiabetic mice from VREF122 induced lethal bacteremia. All live mice remained healthy for additional 30 days of observation when the experiment was terminated. The phage dose effect on the state of health of the infected animals was visible, as shown in Fig.5a and b. The survival of the mice was observed for one month.

### Study of the immune response against ϕ GACP in diabetic and non diabetic mice

The blood was collected from the optical vein of mice, and serum was separated from both IgG and IgM titers of anti-phage$ GACPantibodies were detected subsequently with 3000-fold and 500-fold, respectively groups. The anaphylactic reactions that appeared are adverse and detected no significant differences in core body temperatures or other disastrous events.

## Discussion

The bacteriophages are highly specific for the target bacteria without affecting the normal micro-flora, are effective against multiple drug resistant bacteria. However, phages were discovered nearly a century ago **[21].** Western medicine’s interest in them as therapeutic agents was relatively short-lived because of the eventual discovery and instant success of antibiotics and in part because of the highly empirical and counterproductive approach used by phage practitioners in the early era. In the modern era 1980s and 1990s, some rigorously controlled animal experiments have been conducted animal experiments have been **[7, 22]**, .but the clinical reports in this same era have been subjective rather than unfolding controlled studies **[23].**

The present study emphasized the isolation of MDR strains of *E.faecalis,* inoculated with sewage samples obtained from different local drainages, and phage were harvested and then subjected to the spot test. In the spot test, phage formed a clear zone on *E.faecalis* VREF 122. The results possessed a lytic nature against isolates of *E.faecalis* 122 as positive. In study conduct, the phage strain isolated ENB6 was found to form plaques on 57% of the VRE clinical isolates, and it inhibited bacterial growth of an additional 22% of the strains, exhibiting an antibacterial effect against 79% of the strains in collected samples **[20]**. Further reports on phage Φ SH-56 forming plaques on more than 79% of the MDR enterococci isolates tested and inhibits 7% growth, thus providing an antibacterial effect against more than 86% of approximately 32 clinical isolates of MDR enterococci isolated from diabetic foot cases has been reported **[24].**

In a study, the one-step growth curve is carried out to identify the different phases of the phage infection process. The phage Φ AB2 adsorbed efficiently had a short latent period and caused complete lysis. Upon infection of *A. baumannii* ATCC 17978 in the early-exponential phase (OD_600_=0.6) with Φ AB2 at an MOI of 0.001, cultures were cleared 2×10^10^ PFU/ml of progeny was obtained **[25].** The adsorption rates Φ AB2 to the *A. baumannii* ATCC 17978 cells showed approximately 75 % of phage particles absorbed to host cells within 2 mins, 95 % in 4 min, and nearly 100% in 10 mins. In the tube lysis test, the stool phages JS4 and JS94.1, the environmental water phages JSD.1, and JSL.6 showed a complementary lytic potential of pathogenic *E. coli* strain collected. The tested K803 strain, progeny phage was detected in broth culture infections at about 40 min post-infection, and phage titer increased later, sometimes in a biphasic way **[26].**

Our study of phage ϕ GACP has shown the biological parameters’ measurements as adsorption rate, latent period, and burst size. Comparing these parameters of ϕ GACP, the adsorption rate within 5 min was about 90%, the latent period was about 25min and the burst size was about 110–120 and an average burst size 5×10^8^ PFU/ml. As medians of latent period and burst size in tailed phage are typically 40–60min and 50–100, respectively and was capable of infecting a broad spectrum of *E. faecalis* strains, causing completed and confluent lysis, indicating that ϕ GACP as a therapeutic agent to control infections caused by MDR *E.faecalis.A* study on phage 4D against *E. faecalis* showedlatent period, and burst size for tailed phages is usually 40–60 min and 50–100. A one-step growth curve for ϕ4D showed a latent period (defined as the time interval between the adsorption and the beginning of the first burst) of about 25 min and an average burst size of about 36 PFU/cell **[27].**

Transmission electron microscopy (TEM) of phage ϕ4D revealed an isometric head and a contractile tail with a baseplate. The tubular rear was 164 ± 4.5 nm in length, and the head was 74 ± 4 nm in diameter **[27].** According to Ackermann’s classification, ϕ4D was assigned to the Myoviridae family based on these morphological characteristics. A study on *E. faecalis* ϕSH-56 confirmed the presence of an icosahedral head, about 65 nm in diameter, and a 100 nm long tail, morphologically like phages belonging to Siphoviridae **[23]**. This study of $ GACP reveals that it possesses isometric heads of 100 nm in diameter and conspicuous capsomers, striated 140 long tails, a double base plate, and globular structures at the tail tip of family myoviridae, subfamily Spounavirine family. Phage ENB6 of vancomycin-resistant *E.faecium* has shown a moderately elongated head, 99.7 nm long and 84.4 nm wide, and a contractile tail (199.1 nm long). The phage is of the A2morphotype **[28]**, as observed by negative staining with uranyl acetate.

Many sepsis mouse models are typically used to assess phage therapy against nosocomial bacteria **[20, 28, 24, 29]**. In a study to examine therapeutic effectiveness and the effect of host sensitivity difference on the phage *in-vivo,E. faecalis* sepsis mouse models using BALB/c mice were set up using two strains, EF14 and VRE2. EF14 had phage sensitivity about 32 times greater than that of VRE2 (efficiency of plating [EOP] against EF14, 1; EOP against VRE2, 0.032)**[30]**. After intraperitoneal bacterial inoculation at different concentrations to mice, the minimum lethal bacterial dosage of EF14 and VRE2 was determined to be 1.0 x 10^10^ and 4.2 x 10^9^ bacteria; these dosages resulted in 100% lethality within 2 days. These bacterial dosages were used for our experiments to assess mouse rescue by phage. In diabetic mice, *S.aureus* infection with multiple doses of STZ [ 120 mg (kg body weight)^−1^] is induced in the mouse model. STZ and fasting blood glucose levels after STZ treatment animals with blood glucose levels > 250 mg/dl were considered a diabetic model **[31, 32]**. The STZ diabetic model may serve as a better model since diabetes can be initiated at a younger age when all animals have reached maturity and with negligible weight loss **[33, 34].**

In our animal experiment, the mice were induced with 180mg/dl with STZ to make diabetic, and the glucose level is maintained in the induced mice at 300mg/dl. The lethal dose of *E.faecalis* VREF122 to mice was determined by injecting 3×10^8^, diabetic and non-diabetic mice. In the study reported by Biswas et al., vancomycin resistant *E. faecium* bacteremia with MLD 10^9^ CFU when inoculated was found to be fatal within 48 h **[20]**, thus 3×10 ^9^ CFU as used for the further experiment in our study.

A study on therapeutic effectiveness in vivo of Φ EF24C observed no abnormal mouse behavior or altered survival rate following administration of saline or phage alone (1.0 x10 ^12^ PFU); thus, the phage rescue experiments were considered to be conducted without bias **[30].** The dose-dependent effectiveness is observed to be less than MOI of 0.01 and 0.1 of EF14 and VRE2 mouse sepsis models, respectively, and was similar to results of Φ EF24C efficiently rescuing mice infected with both EF14 and VRE2 at an MOI of 0.01. In the present study single dose of ϕ GACP 3×10^10^ PFU was applied intraperitoneally (i.p) after 20 min VREF 122 challenge and showed significant rescue of 100% in both diabetic and non-diabetic mice from lethal bacteremia. Biswas et al. showed the phage ENB6 9×10^9^ single dose was administrated i.p 45 min after the challenge with the MLD of bacteria. By 24 h dose effect on the state of health of the infected animals was visible. At higher doses of phage multiplicities of infection of 0.3 and 3.0, 100% of the animals survived, and only minimal signs of illness were observed **[20]**.

To analyzed phage treatment safety with the chemotherapeutic treatment of diabetic and non-diabetic single of single i.p dose of vancomycin [2mg (Kg body weight)’ ^−1^ showed 70% percent protection in both groups.If there is an antagonistic effect, the mice no demonstrated changes with the comparable grouping of phage and antibiotic induction. A comparative investigation has been reported **[31]**. These outcomes additionally concur with prior studies **[17, 35, 36]**, in which a stamped distinction in the impact of phage treatment was in bunches rewarded with bacteriophages contrasted with antibiotics agents.Anyway, no investigations have been accounted for *E. feacalis* in induced and non induced diabetic mice.

There is an accord that the general condition of the Slopek and Kucharewica-Krukowska **[20]** animal or patient at the beginning of phage therapy affects the patient’s immunity. In the present study, IgG and IgM were estimated in diabetic and non-diabetic following 28 days of single dose injection of ϕ GACP, increased in the titer background by 3000 fold and 500 fold respectively in both groups. No critical distinction was found among diabetic and non-diabetic IgG and IgM titers against ϕ GACP. No anaphylactic reactions, changes in core body temperature, or other adverse events were observed in the two groups. In an immune response study of phage ENB6 against CRMEN 44 VRE strain, IgG and IgM levels raised against the phage expanded above 3,800-fold and 5-fold, respectively. Anaphylactic reactions were negative, and no changes in core body temperature or other adverse events were observed in the mice for the multiple injections of phage **[20].**Similar findings were reported for patients with severe combined immune deficiency (SCID) described by poor or undeveloped detectable antibody responses to phages even after repeated injections **[18, 37]**.

